# *Cis*-regulatory Elements and Human Evolution

**DOI:** 10.1101/005652

**Authors:** Adam Siepel, Leonardo Arbiza

## Abstract

Modification of gene regulation has long been considered an important force in human evolution, particularly through changes to *cis*-regulatory elements (CREs) that function in transcriptional regulation. For decades, however, the study of *cis*-regulatory evolution was severely limited by the available data. New data sets describing the locations of CREs and genetic variation within and between species have now made it possible to study CRE evolution much more directly on a genome-wide scale. Here, we review recent research on the evolution of CREs in humans based on large-scale genomic data sets. We consider inferences based on primate divergence, human polymorphism, and combinations of divergence and polymorphism. We then consider “new frontiers” in this field stemming from recent research on transcriptional regulation.

## Introduction

The chimpanzee has long presented a conundrum for human geneticists. The orthologous proteins of humans and chimpanzees are more than 99.5% identical [1], yet the two species differ profoundly across a broad spectrum of apparently unrelated phenotypes. This evident paradox led King and Wilson to speculate, famously, that differences in gene regulation, rather than protein-coding sequences, might primarily explain differences in physiology and behavior between humans and chimpanzees [2] (see also [3, 4]). This proposal—while bold—in a sense grew naturally out of Jacob and Monod's research over a decade earlier establishing that the “program” for gene regulation was, in large part, written in DNA [5]. For, as Jacob and Monod themselves recognized [6], if regulatory programs were encoded in the genome, then they were subject to modification by mutation and natural selection, just as protein structure was.

These early conjectures about regulatory evolution were alluring, but for a long time they remained frustratingly abstract and unsubstantiated. In those days, few details could be provided about precisely which regulatory sequences changed, how much, and with what effect. During the ensuing decades, however, indirect evidence and anecdotal examples began to accumulate in support of the idea that *cis*-regulatory elements (CREs) associated with transcriptional regulation played a particularly central role in regulatory evolution [7–9]. (For the purposes of this article, CREs are regulatory sequences relatively near their target gene, typically no more than about a megabase from the transcription unit; we will focus on CREs involved in transcription.) Nevertheless, direct, large-scale support for the prominence of CREs in the evolution of form and function was lacking, and these claims remained controversial [10].

During the past few years, it has finally become possible to examine the evolution of CREs directly on a genome-wide scale, owing to the availability of genomic data describing both genetic variation and regulatory elements. This review will cover major developments over the past decade in the study of human CREs and their role in human evolution, with a particular focus on studies that have leveraged the large public data sets released over the past 2–3 years. Along the way, we will discuss various challenges that arise in the interpretation of these data sets. We will end with a brief survey of new developments in the study of transcriptional regulation that have the potential to enrich studies of human evolution.

## The Old Wave: Studies Based on Interspecies Divergence

A central principle of molecular evolution holds that inferences about natural selection can be made by comparing rates of nucleotide substitution in sites of functional importance with those at sites expected to have little or no influence on fitness. This principle is based on the expectation that mutations will occur at approximately equal rates in both functional and nonfunctional sites, but natural selection will alter the rates at which derived alleles reach fixation in functional sites (Figure 1). This idea has been applied for decades to protein-coding sequences, where amino acid altering (nonsynonymous) and non-altering (synonymous) substitutions provide convenenient classes to contrast [11–13].

**Figure 1:**
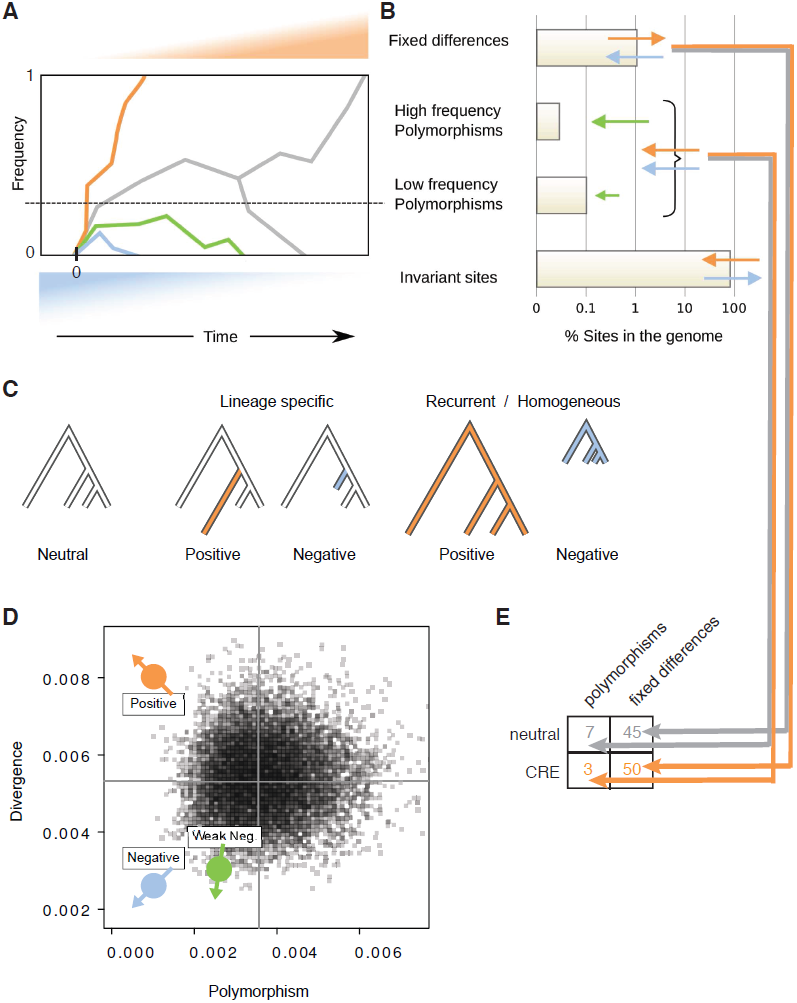
(A) Frequency as a function of time for hypothetical mutations experiencing neutral drift (gray), weak negative (green), strong negative (blue), or positive (orange) selection. The plot assumes a new mutation occurs in a single individual in the population at time 0. Neutral drift typically causes mutations to be lost (lower gray fork) but occasionally drives them to fixation (upper gray fork). Negative selection essentially guarantees eventual loss, but if it is sufficiently weak (green plot), mutations may segregate at low frequencies for some time. Positive selection (orange plot) causes mutations to reach fixation at higher rates than neutral drift. Notice that the time until fixation or loss is substantially reduced for mutations under strong selection (positive or negative), implying that they are unlikely to be observed in a polymorphic state. (B) Steady-state numbers of invariant sites, low frequency (derived allele) polymorphisms, high frequency polymorphisms, and fixed differences under neutral drift, expressed as hypothetical percentages of nucleotide sites. These represent equilibrium frequencies for the process depicted in panel (A) for a given divergence time, assuming a steady flow of new mutations. Positive selection (orange arrows) increases fixed differences, reduces invariant sites, and reduces polymorphisms. Strong negative selection (blue arrows) reduces fixed differences and polymorphisms and increases invariant sites. Weak negative selection (green arrows) is similar but allows some low frequency polymorphisms to remain. (C) Phylogenies with branch lengths proportional to rates at which fixed differences occur along lineages. Positive or negative selection can be identified by significant increases or decreases, respectively, in the fixation rates relative to the neutral expectation. Different likelihood ratio tests can identify lineage-specific or recurrent/homogeneous selective pressures. (D) Scatter plot of polymorphism vs. divergence rates under neutral drift, generated by simulations based on parameters reflecting real human populations [44] (black points). Colored points show hypothetical positions of sequences under positive (orange), strong negative (blue), and weak negative (green) selection. Notice that positive and negative selection are distinguishable by their joint effects on polymorphism and divergence rates, but not by polymorphism rates alone. (E) 2 × 2 contingency table used for McDonald-Kreitman (MK) test for selection on a *cis*-regulatory element (CRE). The test evaluates the probability of the observed data under the null hypothesis that the relative polymorphism and divergence counts are independent of the labels “neutral” and “CRE”. The classes of sites are chosen to be similar to one another to avoid potential biases from mutation rate variation and demography. Rejection of the null hypothesis therefore implies a departure from the neutral expectation of equal fixation rates. Note the connections with the visual representations used in panels (B) and (D). The MK test can be thought of as comparing the relative heights of the first bar and the next two bars combined in panel (B), for neutral vs. CRE sites (see arrows). It can also be thought of as testing for extreme departures from a diagonal line in panel (D) running through the neutral points from bottom left to top right. In this case, the counts reflect an excess of fixed differences in the CRE, suggesting positive selection. Notice that strong negative selection is not a problem for the MK test, because it reduces the effective mutation rate, but weak negative selection can bias the test by partially canceling the effects of positive selection.

The sequencing of the chimpanzee genome [1] enabled analogous methods to be applied genome-wide to putative CREs in hominids. For example, Keightley et al. examined sequences in upstream regions and first introns of genes and contrasted them with other intronic sequences assumed to be neutrally evolving [14]. They found that putative regulatory sequences showed almost no evidence of constraint in hominids, but were significantly constrained in mouse and rat. Finding no signs of positive selection, they argued that regulatory sequences in hominids had experienced “widespread degradation” due to their reduced effective population sizes (see also [1, 15, 16]). Soon afterward, Khaitovich et al. analyzed human-chimpanzee divergence patterns in promoter regions together with data on mRNA expression. Interestingly, they found that human-chimpanzee divergence in gene expression (normalized for intraspecies diversity) was much more pronounced in the testis than in the brain or several other tissues, possibly reflecting positive selection due to differences in mating strategies. They did find an excess of lineage-specific changes in expression of brain genes in human relative to chimpanzee.

Haygood and colleagues improved on the statistical methology of previous studies by developing a phylogenetic likelihood ratio test analogous to those used for protein-coding sequences [17, 18] for lineage-specific elevations in substitution rates in promoter regions [19] (see Figure 1C). Based on alignments of the human, chimpanzee, and rhesus macaque genome sequences, Haygood et al. found evidence of positive selection acting on the promoters of at least 250 genes. High-scoring genes were significantly enriched for roles in neural development and function, nutrition, and metabolism, suggesting an important role for CREs in human cognitive, behavioral, and dietary adaptations. Another series of studies, based on similar statistical methods, tested conserved noncoding sequences for “accelerated” evolution in humans [20–23].

The first large-scale study of primate evolution to make use of newly emerging chromatin immunoprecipitation and microarray (ChIP-chip) data for TF binding was carried out by Gaffney and colleagues [24]. The authors collected ChIP-chip data from seven previously published studies, and then analyzed patterns of divergence at bound sites in the human, chimpanzee, and rhesus macaque genomes, comparing the regulatory sequences with “control” regions. They also considered transcription factor binding sites (TFBSs) recorded in the TRANSFAC database. Using a simple divergence-based estimator, they predicted that about 37% of mutations in TFBSs were deleterious, about half the fraction estimated for 0-fold nonsynonymous sites in coding sequences.

## The New Wave: Studies Based on Intraspecies Polymorphism

Divergence-based analyses, while informative, are fundamentally limited by the relatively long evolutionary time periods associated with the accumulation of fixed differences between species. Irregularities in the evolutionary process during these periods—for example, due to changes in the locations or boundaries of CREs, or changes in selective pressures—can weaken the signal of natural selection, causing its influence to be underestimated. This problem can be mitigated by working instead with data describing genetic variation within a single species [25]. Intraspecies polymorphism provides a window into much more recent evolutionary processes, on the time scale of genealogies of individuals rather than species phylogenies (for humans, roughly 1M years or less), during which the evolutionary process is likely to be more homogeneous. It has been demonstrated at numerous individual loci that patterns of human polymorphism can reveal the influence of natural selection on CREs [26–29].

Several groups have recently used this approach in genome-wide analyses of CREs, taking advantage of the abundant high-quality human polymorphism data now available. Because polymorphisms are sparse along the genome, these groups have generally pooled data across many similar loci. For example, Mu and colleagues examined human polymorphism data from the 1000 Genomes Project in various classes of coding and noncoding elements, including ChIP-seq-supported TFBSs [30]. The authors found that TFBSs were significantly constrained, but less so than coding sequences. Negative selection dominated in their tests, with no sign of pervasive positive selection. They observed stronger constraint in bound than in unbound TFBSs, in TFBSs proximal to transcription start sites (TSSs) than in ones distal to TSSs, and in TFBSs with strong rather than weak ChIP-seq signals. The related work of Khurana et al. further showed that mutations that decrease the matching score of a motif were enriched for rare alleles compared to ones that did not [31]. However, Khurana and colleagues found evidence of contributions from positive selection as well as negative selection in several types of regulatory elements, including DNase-I hypersensitive sites (DHSs) and sequence-specific TFBSs.

In another analysis of 1000 Genomes data, Ward and Kellis examined mean SNP density, heterozygosity, and derived allele frequency in various noncoding regions identified as having “biochemical activity” by the Encyclopedia of DNA Elements (ENCODE) project [32]. They observed significant constraint in putative regulatory regions identified by a wide variety of experimental assays. Interestingly, they found such evidence both for regions that were conserved across mammalian species and ones that were nonconserved, suggesting that a substantial fraction of functional noncoding elements reside outside of mammalian-conserved regions. In a similar study, Vernot et al. analyzed 53 high coverage individual genome sequences in more than 700 motifs within DHSs from 138 cell and tissue types, finding that many of these motifs were signficantly constrained [33].

A separate line of research has considered patterns of nucleotide diversity in flanking sequences of noncoding regions conserved across mammals, which are likely enriched for CREs [34–37]. These studies have come to conflicting conclusions, with some arguing for a prominent role for hitchhiking (HH) from positively selected sites in regulatory elements [34, 37], and others maintaining that the observed patterns are more consistent with background selection (BGS) from negative selection [35, 36]. More work is needed to resolve this controversy over the relative roles of positive and negative selection in shaping CREs.

## A Fusion of the Old and the New: Joint Consideration of Divergence and Polymorphism

Population genomic data, too, has limitations when used as the sole source of information about natural selection. As noted above, it can be difficult to distinguish between positive and negative selection based on patterns of polymorphism alone (both forces reduce diversity; see Figure 1). Another major challenge is accounting for the effects of population bottlenecks, expansions, and other demographic processes, which can profoundly influence allele frequencies even in the absence of natural selection [38]. These problems can be alleviated by jointly considering intraspecies polymorphism and divergence from a neighboring species, an idea that has been used for decades in the analysis of protein-coding genes [39–41]. Classical approaches of this kind, such as the McDonald-Kreitman (MK) test [40], compare relative rates of polymorphism and divergence in putatively functional and nonfunctional (typically, nonsynonymous and synonymous) classes of sites. Under neutral drift, fixation should occur randomly for both classes of sites, causing the ratios of polymorphisms and fixed differences to be approximately equal. Departures from this neutral expectation provide information about natural selection (Figure 1).

An early attempt at a joint analysis of polymorphism and divergence of CREs, by Torgerson and colleagues, examined conserved noncoding regions flanking more than 15,000 protein-coding genes, using polymorphism data from 15 African Americans and 20 European Americans as well as the chimpanzee genome [42]. The authors made use of an extension of the MK test that permits estimation of selection coefficients [43], adapting it for use with noncoding sequences. Consistent with previous analyses, they found clear evidence of purifying selection in these regions. In addition, they found a significant excess of fixed differences relative to polymorphic sites, indicating positive selection on at least some CREs. In the study discussed above [24], Gaffney and colleagues also made limited use of polymorphism data, attempting to compute the fraction of fixed differences driven by positive selection (*α*) in CREs using a simple estimator based on the MK framework (see [41]). In contrast to Torgerson et al., they found no significant evidence of positive selection on CREs, but their power appeared to be quite weak.

Arbiza and colleagues attempted to address previous limitations in both models and data in a large-scale analysis of TFBSs based on ChIP-seq data from the ENCODE project [44]. Using a new probabilistic model and inference method called INSIGHT, the authors analyzed 1.4 million binding sites from 78 TFs, together with genetic variation data from the human, chimpanzee, orangutan, and rhesus macaque genome sequences, and 54 high-coverage human genome sequences. They found strong evidence of both positive and negative selection in TFBSs, with somewhat more positive selection, more weak negative selection, and less strong negative selection than in protein-coding genes. The authors estimated that, overall, there have been at least as many adaptive substitutions in CREs as in protein-coding genes since the human-chimpanzee divergence, consistent with King and Wilson's conjecture almost forty years earlier.

Another interesting observation from this study was that regulatory regions exhibited a large excess of weakly deleterious segregating mutations compared with protein-coding genes, suggesting considerable genetic load associated with gene regulation. This finding is concordant with a recent analysis of genetic association data, which found that regulation-associated DNase-I hypersensitivity sites accounted for almost 80% of the heritability for 11 common diseases [45]. Together, these findings suggest that a shift toward weaker negative selection in CREs may somewhat paradoxically result in an enrichment for heritable disease-causing segregating variants, because these variants are less efficiently eliminated by natural selection than those in protein-coding genes.

## The Next Frontier

Most studies of *cis*-regulatory evolution in humans, including all of those discussed so far, have assumed that binding sites maintain stable positions at orthologous genomic locations over evolutionary time, and that fitness effects can be measured by patterns of variation at individual nucleotide positions. In reality, however, natural selection acts on nucleotides in TFBSs only indirectly, through the effects of those nucleotides on transcriptional output. These effects, in turn, occur through a complex and incompletely understood set of physical interactions involving multiple TFs and cofactors, the core transcriptional machinery, the DNA sequence, the local chromatin, and the surrounding aqueous environment [46, 47] (see Figure 2). Recognizing the full complexity of transcriptional regulation will be essential for a complete understanding of its evolution in humans and other species.

**Figure 2:**
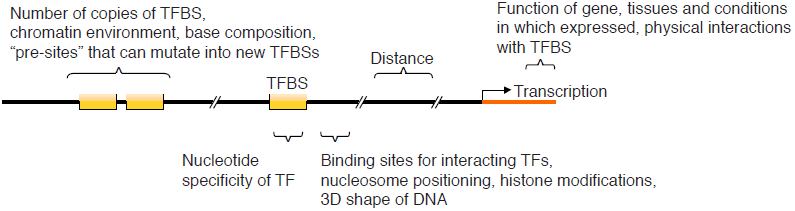
Some of the many factors that may influence the evolution of *cis*-regulatory elements.

### Biophysical Models of Binding-Site Evolution

A pioneering series of papers by Lässig and colleagues began to explore this complex intersection of biophysics and evolution using models that treated the free energy of TF binding to DNA as a quantitative phenotype, which served as the basis of an explicit fitness landscape. Evolutionary trajectories over this landscape were then considered [48–51] (see also [52]). Despite assuming an additive model for nucleotide-specific binding energies, the authors obtained highly nonlinear fitness landscapes, reflecting epistasis between regulatory nucleotides. In both prokaryotes and yeast, they found evidence for widespread compensatory mutations and relatively frequent gain and loss of binding sites.

Following these observations, Moses developed statistical tests for natural selection in terms of changes in predicted binding affinity resulting from single nucleotide changes under standard position-weight-matrix (PWM) models of binding [53]. Another study showed that evolutionary events tended to preserve binding affinity in Drosophila [54]. More recently, Bullaughey studied the evolution of enhancers by combining a thermodynamic sequence-to-expression model [55] with a Gaussian expression-to-fitness model [56]. His simulation study suggested strong interdependencies between nucleotides and an important role for neutral substitutions in changes to the functional organization of enhancers. Finally, in an analysis of well characterized *cis*-regulatory modules in Drosophila, He et al. found bulk evidence for positive selection contributing to both gain and loss of binding sites and for purifying selection maintaining existing TFBSs [57].

Another recent series of papers has focused on the development of improved biophysical models of TF binding to DNA, generally without consideration of evolution. A full review of this literature is outside the scope of the present article, but examples include models that consider combinatorial interactions among TFs [58–62], nucleosome positioning and/or chromatin accessibility [63–66], and the three-dimensional structure of DNA binding sites [67] (see [47] for a related review). More work is needed to consider the evolutionary implications of biophysical models of this type, but it seems likely that inferences of the distribution of fitness effects of regulatory mutations in humans will change significantly when richer, more realistic models of binding site structure and function are considered.

### Improved Characterizations of Binding Affinity

Even the sophisticated biophysical models discussed in the previous section have tended to maintain the assumption of additive contributions of individual nucleotides to TF binding affinity, corresponding to an assumption of site independence in statistical motif models [68, 69]. This assumption appears to be adequate for most TFs, but numerous violations have been observed [70–72]. Nevertheless, statistical methods that attempt to recover the full correlation structure of TF binding preferences from sequences [71, 73, 74] have not been widely adopted.

These challenges have led to intense interest in harnessing high-throughput genomic technologies to produce direct measurements of binding affinity for all possible binding sites and large numbers of TFs. Widely variable strategies have been employed, including microwell-based assays [75, 76], protein-binding microarrays [67, 77–79], mechanically induced trapping of molecular interactions (MITOMI) [80], high-throughput systematic evolution of ligands by exponential enrichment (SELEX), [81–83], and, most recently, adaptation of the Illumina sequencing platform to directly measure binding affinities of proteins to DNA [84] (see [85] for a review as of 2010). In addition to finding further evidence of positional inter-dependence [79, 83, 84, 86, 87], studies based on these techniques have revealed, among other features, unexpected dimeric modes of binding [82], numerous TFs that recognize multiple sequence motifs [79], and important influences of sequences flanking core binding sites owing to their effects on DNA shape [67, 83]. However, the rich models of binding affinity enabled by these powerful technologies have yet to be integrated into evolutionary models.

### Evolutionary Turnover of *Cis*-Regulatory Elements

As alluded to in the previous section, there is strong evidence that individual CREs in many species, including humans, are gained and lost over time, a phenomenon known generally as “turnover” [88–90]. Turnover of CREs has been extensively studied over the past decade [56, 57, 91–98] but, overall, it remains poorly understood. For example, it is still unclear how frequently turnover occurs overall, how much it varies across species, TFs, and genomic contexts, how commonly gains and losses are compensatory, and how all of these processes impact inferences of selection. Recent studies that make use of high-throughput functional genomic techniques applied uniformly across species [99, 100] have helped to shed additional light on turnover of CREs, but these studies also have limitations. For example, it is not clear how many of the assayed binding events directly influence gene expression, what role false negatives and false positives play in apparent differences, and in some cases sample sizes have been insufficient to distinguish within-species variation from between-species divergence. In our view, it will be essential to develop improved methods for integrating evolutionary and biophysical models with large-scale functional genomic data, to develop a more complete understanding of the complex processes by which CREs evolve.

## Acknowledgments

A.S. is supported by NIH grants R01-GM102192 and R01-HG007070. L.A. is supported by NIH grant R01-HG006849 to Alon Keinan. The authors’ thinking about transcriptional regulation has benefited from many valuable discussions with Andre Martins, Charles Danko, Leighton Core, and John Lis.

